# Modulation of Individual Alpha Frequency with tACS shifts Time Perception

**DOI:** 10.1101/2020.07.23.218230

**Authors:** G. Mioni, A. Shelp, C. T. Stanfield-Wiswell, K. A. Gladhill, F. Bader, M. Wiener

## Abstract

Previous studies have linked brain oscillation and timing, with evidence suggesting that alpha oscillations (10Hz) may serve as a “sample rate” for the visual system. However, direct manipulation of alpha oscillations and time perception has not yet been demonstrated. Eighteen subjects performed a time generalization task with visual stimuli. Participants first learned the standard intervals (600 ms) and then were required to judge the new temporal intervals if they were equal or different compared to the standard. Additionally, we had previously recorded resting-state EEG from each subject and calculated their Individual Alpha Frequency (IAF), estimated as the peak frequency from the mean spectrum over posterior electrodes between 8 and 13 Hz. After learning the standard interval, participants performed the time generalization task while receiving occipital transcranial Alternating Current Stimulation (tACS). Crucially, for each subject, tACS was administered at their IAF or at off-peak alpha frequencies (IAF±2 Hz). Results demonstrated a linear shift in the psychometric function indicating a modification of perceived duration, such that progressively “faster” alpha stimulation led to longer perceived intervals. These results provide the first evidence that direct manipulations of alpha oscillations can shift perceived time in a manner consistent with a clock speed effect.

## Introduction

Despite temporal processing having a fundamental role in daily living, the neural mechanisms underlying time perception are still being uncovered. Recent research in this area has begun focus on the role of endogenous oscillations in temporal perception and action. Yet, the role of any particular neural oscillation in temporal perception remains to be determined (Wiener & Kanai, 2016). In search for an internal timekeeper, some theories (Treisman, 1963) have suggested that the rate of an internal pacemaker would be driven by neural oscillations in the alpha range (8– 12Hz). Faster alpha rhythms would result in longer estimates of time than slower alpha rhythms, considering that more pulses would accumulate during the same physical time interval (Treisman, et al., 1990; 1994). Given recent findings that individual alpha frequency (IAF) correlates with temporal illusions (Cecere, Rees, & Romei, 2015), it remains plausible that fluctuations of alpha peaks could modulate perceived duration. However, the relationship between IAF and modification in subjective timing has not been clearly established.

Associations between cognitive performance and endogenous modulations of oscillatory neural activity in the individual alpha frequency (IAF) range have been established in a number of studies (Klimesch, 1999). Indeed, previous research has revealed that IAF predicts performance on a variety of perceptual (e.g., Cecere et al., 2015; Samaha & Postle, 2015) and cognitive (e.g., Bornkessel, et al., 2004; Klimesch, Doppelmayr, & Hanslmayr, 2006) tasks. If these associations are functionally relevant, then it should be possible to influence cortical oscillations with non-invasive brain stimulation techniques (Helfrich, et al. 2014; Ruhnau, et al. 2016) and thereby modulate behavioral performance (Klimesch, 2012). Here, we applied transcranial alternating current stimulation (tACS), under the hypothesis that it can induce predictable changes in oscillatory cortical activity and alter perception in a temporal generalization task.

Previous studies have shown that the phase of alpha oscillations at the onset of two temporally-proximal visual stimuli predicted whether they were perceived as occurring simultaneously or asynchronously (Varela, et al., 1981; Samaha & Postle, 2015) suggesting that alpha cycles reflect sequential discrete “frames” of visual perception (Busch et al. 2009; Busch & van Rullen, 2014). More recently, Cecere and colleagues (2015) observed that stimulus temporal proximity determines whether or not inputs coming from different modalities are bound or not. Using a flash-sound illusion paradigm, in which a single flash bound by two tones produces an illusory second flash, the authors were able to modify the temporal window of integration, indexed by how close in time the tones had to be to induce an illusory percept, by applying tACS during task performance at IAF or IAF ±2Hz. Compared to performance at tACS IAF, the temporal window was wider at IAF-2Hz and narrower at IAF+2Hz, indicating that a faster alpha rate was associated with less susceptibility to the illusion. Similarly, Zhang and colleagues (2019) showed that applying tACS at different frequency in the α-band changed subjects’ perceptual switch rate in the binding of bistable color-motion stimuli. Taken together, previous studies provided converging evidences for a causal link of IAF and temporal resolution of visual perception (see also Samaha & Poste, 2015; Samaha, Bauer, Cimaroli, & Poste, 2015).

The present study aims to investigate the link between modification in IAF and variation in perceived duration. We first recorded EEG during a 5-min resting state, which was used to extract IAF; then, participants performed a sub-second visual time generalization task while receiving tACS either at their IAF, or at individually accelerated or decelerated rates (IAF±2 Hz). We hypothesized that driving IAF toward slower versus faster oscillations should result in under- or over-temporal estimation, respectively (Figure 1). As an alternative hypothesis, we note that altering IAF may not shift perceptual duration *per se*, but instead alter the resolution at which time intervals are perceived. Under this instance, no biases should be observed, but instead a sharpening/blunting of temporal estimates, leading to differences in the precision of generalization gradients across conditions.

**Figure 1.**
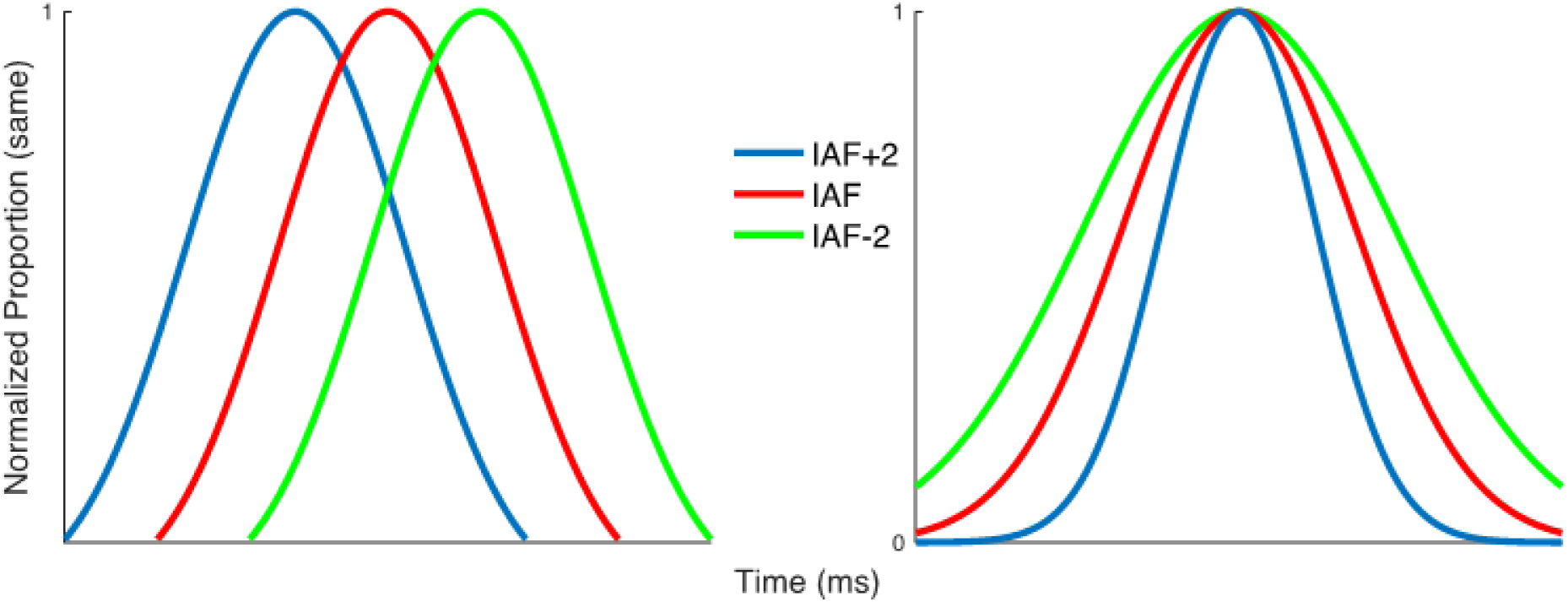
Two possible hypotheses for the manipulation of IAF and time estimation. Left: if IAF indexes the rate of temporal estimation, akin to altering the speed of an internal pacemaker, then increasing/decreasing the speed of this pacemaker would shift temporal estimates, such that a faster/slower IAF leads to intervals perceived as longer/shorter. Right: if IAF indexes the sample rate or resolution of the temporal estimator, then increasing increasing/decreasing the resolution would sharpen temporal estimates, such that a faster/slower IAF leads to more/less precise estimates of duration.

## Materials and Methods

### Participants

Thirty-two right-handed, healthy volunteers (mean age = 22.95, SD = 4.64; range 19-36 years) who met inclusion criteria for brain stimulation were initially contacted for participation in the experiment. Of those contacted, 19 responded and agreed to participate (mean age = 23.15, SD = 9.80); among these one was excluded due to a misunderstanding of the task resulting in uninterpretable data. The final sample included 18 right-handed healthy volunteers (9 males, 9 females; mean age, 22.89, SD = 4.47 years; range, 19–36 years) who met inclusion criteria for brain stimulation participated in the experiment. All subjects gave their informed consent as approved by the Institutional Review Board of George Mason University.

### Procedure

Participants were tested in two separate experimental sessions on separate days. One the first day, participants underwent 5-minute EEG recording and we calculated IAF. On the second testing day, in three separate blocks, participants received tACS over occipital cortex to modulate occipital alpha oscillations at their IAF or at slower (IAF -2Hz) or faster (IAF +2Hz) rates while they were performing the time generalization task, stimulation level was counterbalanced between participants. The tACS blocks were 30 min apart from each other (Figure 2, Top).

**Figure 2.**
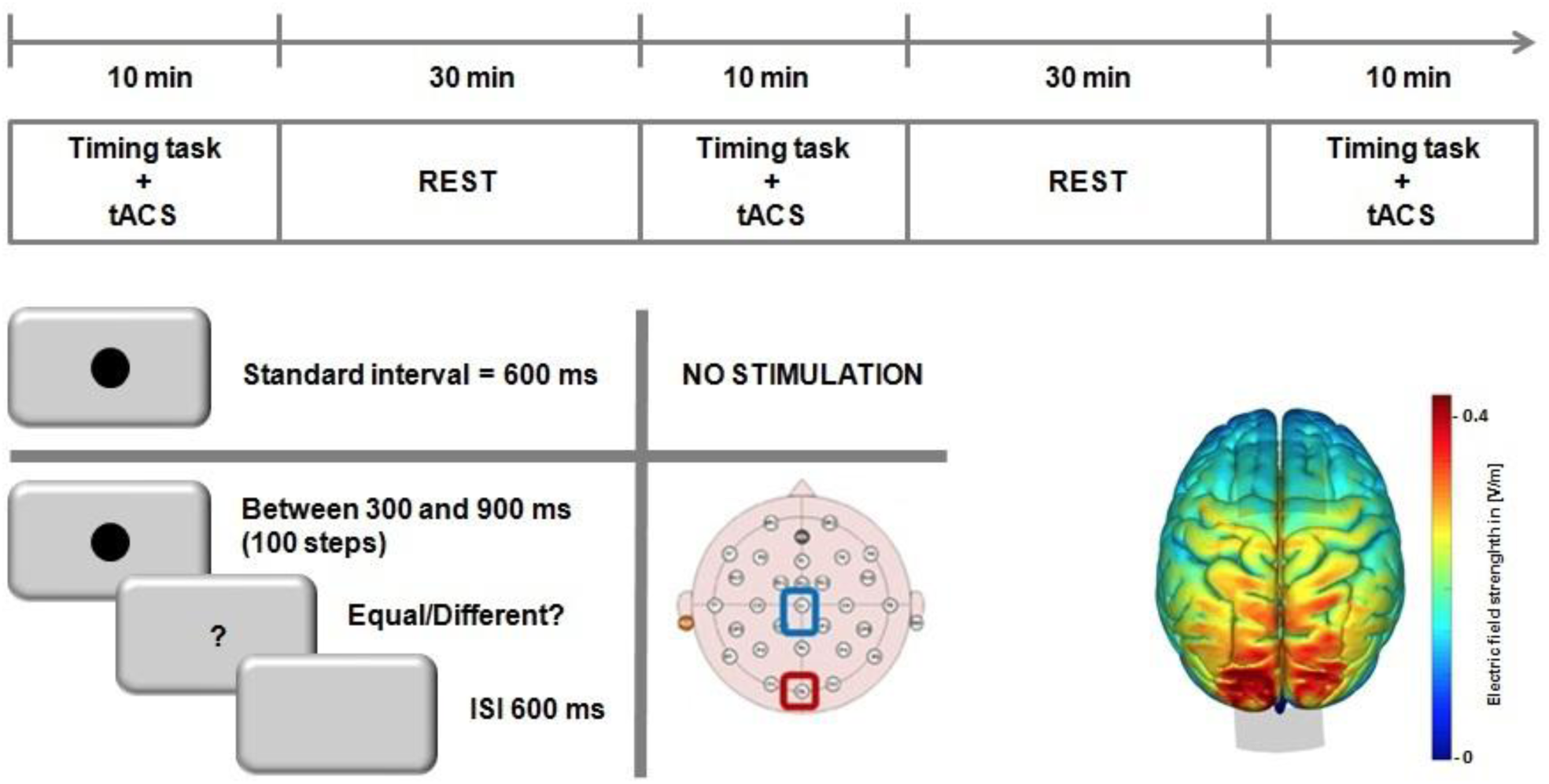
Graphical representation of experimental procedure. Top: timeline of the experiment for the tACS sessions; stimulation levels were counterbalanced across subjects/blocks. Bottom Left: schematic for the temporal generalization task, in which subjects first learned a standard 600ms interval, prior to stimulation, and were then tested on a range of linearly-spaced intervals around the standard. Bottom Right: a simulation of the normalized electric field strength of tACS electrodes, run via SimNIBS displaying maximal field strength over occipital cortex.

### Time generalization task

We chose a time generalization task in which the response requirements (same/different) are non-directional in nature, and therefore resistant to response bias over tasks relying on a directional response (i.e. longer/shorter) (Yates et al., 2012). Participants were instructed to judge if a probe stimulus was the same or different than a previously learned standard. Visual stimuli (grey circle) were presented centrally on a computer screen in a lit room while participants were sat on a comfortable chair at ∼60 cm viewing distance. Stimulus presentation and behavioral responses were recorded by a computer using E-Prime software (Psychology Software Tools, Inc. Version 2.0). In the temporal generalization task, participants were instructed to compare the duration of the presented stimulus with a previously learned standard (Droit-Volet, 2002). During an initial learning phase, participants encoded the standard duration (600 ms), presented ten times (Figure 2 Bottom left). In the subsequent testing phase, participants were given 4 blocks of 56 trials without feedback. Each block included 8 initial presentations of the standard duration (600 ms) as a “top-up” phase, and 48 presentations of different intervals (24 shorter: 300, 400, 500 ms and 24 longer: 700, 800, 900 ms). Participants had to judge whether the comparison duration was similar (equal response) or not similar (different response) to the standard duration by pressing the corresponding key. We used a QWERTY keyboard to record responses and response keys were labelled with a yellow sticker for “yes” and green stickers for “no” responses. Participants responded with their index fingers; the yellow sticker was placed over the “S” letter on the keyboard and the green sticker was placed over the “L” letter on the keyboard.

### EEG recording and analysis

We note that these EEG data were collected from several other experiments concurrently being conducted at our institution; as such, two different EEG recording systems were used. Fifteen participants were recorded using a 64-channel active electrode system (BrainProducts, Munich, Germany) with a sample rate of 1000Hz and impedances kept below 20 kΩ. Three participants were recorded using a 40-channel passive electrode system (NeuroScan) with a sample rate of 1000Hz and impedances kept below 5 kΩ. In both recordings, the electrodes were placed according to the 10-20 system; the online reference was placed at FCz and the ground was placed at AFz in the 64 channel system, whereas the online reference was the left mastoid in the 40-channel system. Data were recorded using an online band-pass filter of 0.1 - 100 Hz for the 64-channel system and a high-pass filter of 0.1 Hz for the 40-channel system.

EEG data were preprocessed offline using EEGLAB (version 12.0.2b; Delorme & Makeig, 2004). First, the data were down-sampled to 500Hz and re-referenced to the common average. Then, we performed an automatic rejection of noisy EEG channels on continuous data and confirmed by visual inspection. The noisy channels were interpolated using spherical splines. Next, we applied an automatic artifact rejection procedure on the continuous EEG data using ERPLAB (Lopez-Calderon & Luck, 2014). A Fourier transform was then conducted on all electrodes using the remaining, artifact-free dataset using the EEGLAB function *spectopo* with a frequency factor of 10. Finally, we extracted the IAF by first detrending the spectral data for 1/f noise, and then averaging the maximum spectral peak (8-13Hz) of the following *a-priori* channels: P1, P2, P3, P4, Pz, P5, P6, P7, P8, PO3, PO4, POz, PO7, PO8, O1, Oz, O2 (Corcoran, Alday, Schlesewsky, & Bornkessel-Schlesewsky, 2018) (Figure 2). For the three subjects using the 40-channel system, the same electrodes were used minus those not available. For each subject, the averaged power spectrum was visually inspected for a clear peak around 10 Hz; any subjects exhibiting a lack of a discernable peak were excluded.

### tACS

The tACS was conducted within a single session and the three blocks of stimulation occurred 40 min apart (cf. Cecere et al. 2015). Positions of Oz and Cz were determined using the 10–20 system. tACS was delivered by NeuroConn DC-Stimulator+ through a pair of rubber electrodes enclosed in saline-soaked sponges and fixed on the head by elastic bands. The reference electrode was placed over Cz and the active electrode was placed over the occipital cortex (Oz). The reference electrode (Cz) had a larger size (35 cm^2^) than the active electrode (Oz, 9 cm^2^) to decrease current density delivered over Cz (Cecere et al., 2015). Impedance was maintained within the safety criteria of the tACS device; if they were not met the experimenter added saline solution and tightened the headband to improve electrode contact with the scalp. tACS was delivered at 1.5 mA peak-to-peak; participants were stimulated for a total of 10 minutes in each block. Subjects completed a post-study questionnaire after each session to assess side effects (tingling, itching sensation, burning sensation, pain, and fatigue). All participants were actively encouraged to report any perception of tACS-induced phosphenes throughout the experimental sessions. To avoid phosphenes throughout the experimental sessions participants received 30-sec stimulation at their IAF and IAF±2Hz before performing the time generalization task; if phosphenes were reported, stimulation would be lowered until none were detected. None of the participants reported phosphenes during any of the 30-sec stimulations prior to the testing phase, nor after any of the stimulation sessions, and so the intensity of stimulation was kept at 1.5 mA for all participants.

To validate the likely physiological source of stimulation, we simulated the effect of our electrode montage using the SimNIBS 2.0 Toolbox (http://simnibs.de/). As in our main experiment, the placement of two electrodes located at Oz and Cz, were simulated with a current of 1.5 mA. The normalized electrical field was simulated via a realistic finite element head model (Thielscher et al., 2009). The results of this analysis revealed a centralized electrical field effect centered over the occipital cortex (Figure 2, bottom right).

### Statistical analyses

Performance was analyzed in terms of the Point of Subjective Equality (PSE), calculated as the time value at which subjects were maximally likely to judge the stimulus as equal and the Weber Ratio (WR), calculated as the normalized variability of measurements. Individual PSEs for each of the three stimulation conditions were estimated using adapted Matlab routines for fitting curves to simultaneity judgment tasks (Alcalá-Quintana & García-Pérez, 2013). Briefly, this routine is based on an independent-channels model for stimulus timing judgment tasks proposed by García-Pérez and Alcalá-Quintana (2012a). The advantage of this model over fitting a simple Gaussian curve is that it is a psychophysical model that explicitly accounts for response errors arising from non-specific factors (e.g. lapses of attention, misreports) (Garcia-Perez & Alcalá-Quintana, 2012). The results of this analysis yielded the PSE, and the WR calculated as the peak spread of the fitted curve divided by the PSE.

PSE and WR data were included in separate repeated-measures ANOVAs with *Condition* (IAF, IAF+2Hz and IAF-2Hz) as within-subjects factors. Significant effects were followed up by post-hoc analyses with Bonferroni correction to reduce the Type I error rate, and the effect size was estimated with the partial eta squared index (η^2^_p_). An observed shift of the PSE for the different condition (IAF, IAF+2Hz and IAF-2Hz) can be interpreted as an indicator of differences in these conditions, with smaller bisection point (left shift of the psychometric function) values meaning longer perceived durations. Concerning WR, higher values indicate higher temporal variability.

## Results

Each participant underwent a single, 5-min eyes-open resting-state EEG session before the temporal task and we extracted IAF. For the IAF analysis, we measured a mean IAF of 10.4 Hz ± 1.02, in the approximate middle of the alpha band. The IAF data were further determined to be normally distributed via a Shapiro-Wilk test [*W*(18)=0.95, *p* = 0.422] (Figure 3).

**Figure 3.**
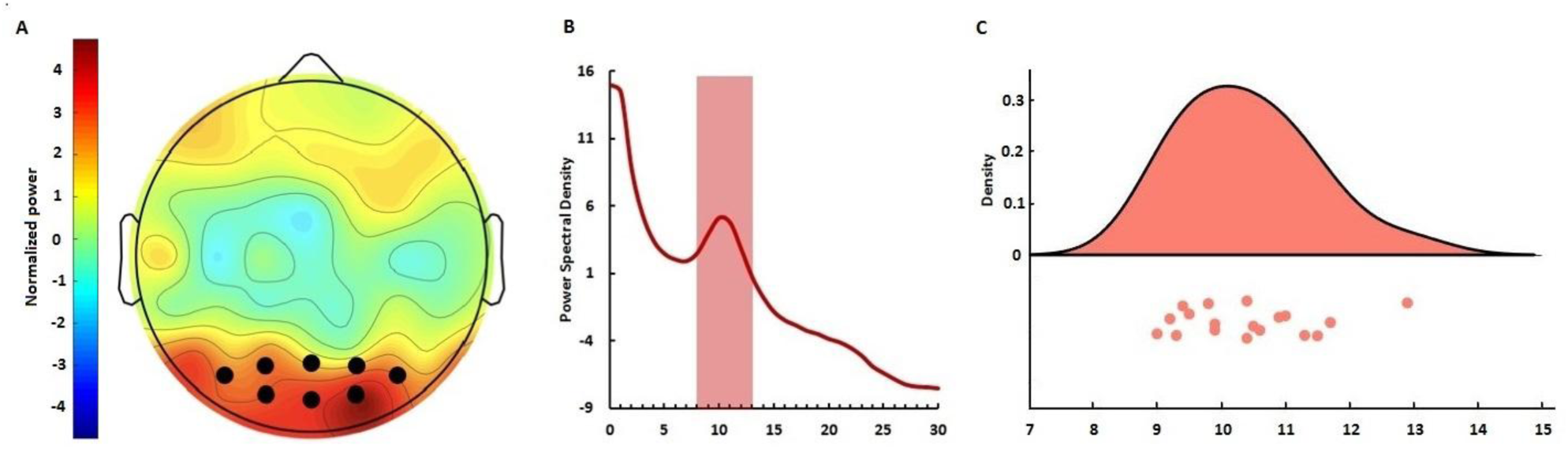
IAF values used for the tACS experiment. **A)** Topographic plot of the average spectral power across subjects in the alpha band. The distribution exhibited strongest power at posterior electrodes; dark circles indicate electrodes used for calculating IAF. **B)** Mean power spectrum (log scale) across the electrodes in (A) exhibiting a characteristic 1/f distribution and typical “peak” in the alpha range. **C)** Raincloud plot (Allen, 2019) of IAF values calculated for each subject; values were normally distributed across the alpha band.

Subjects then performed a temporal generalization task with visual stimuli, in which they were required to judge if a probe stimulus was the same or different than a previously learned, prior to stimulation, standard (600ms). When analyzing the mean proportion of trials on which subjects classified durations as “same”, we initially observed a main effect of tested duration [*F*(6,102) = 15.12, *p* < 0.001, η^2^_p_ = 0.471] and a significant interaction between tACS and duration [*F*(12,204) = 4.613, *p* < 0.001, η^2^_p_ = 0.213] (Figure 4; Figure 5A). Concerning perceived duration (point of subjective equality, PSE), a main effect of tACS [*F*(2,34) = 4.232, *p* = .023, η^2^_p_ = .20] was found; post hoc paired *t*-tests showed significant difference between PSE at IAF-2Hz and PSE at IAF+2Hz [t(17) = 2.869, *p* = .011, Cohen’s *d* = 0.68] but not between PSE at IAF and PSE at IAF+2Hz (*p* = 0.134) and PSE at IAF and PSE at IAF-2Hz (*p* = 0.169) (Figure 5B). Further, a linear contrast effect was observed across PSE values from slower to faster IAF values [*F*(1,17) = 8.234, *p* = 0.011, η^2^_p_ = 0.326], in which the PSE shifted linearly from left-to-right with faster-to-slower tACS rates. The effect of stimulation acted specifically on perceived duration and did not influence temporal variability (Weber ratio) [*F*(2,34) = 1.206, *p* = 0.312, η^2^_p_ = 0.066] (Figure 5C).

**Figure 4.**
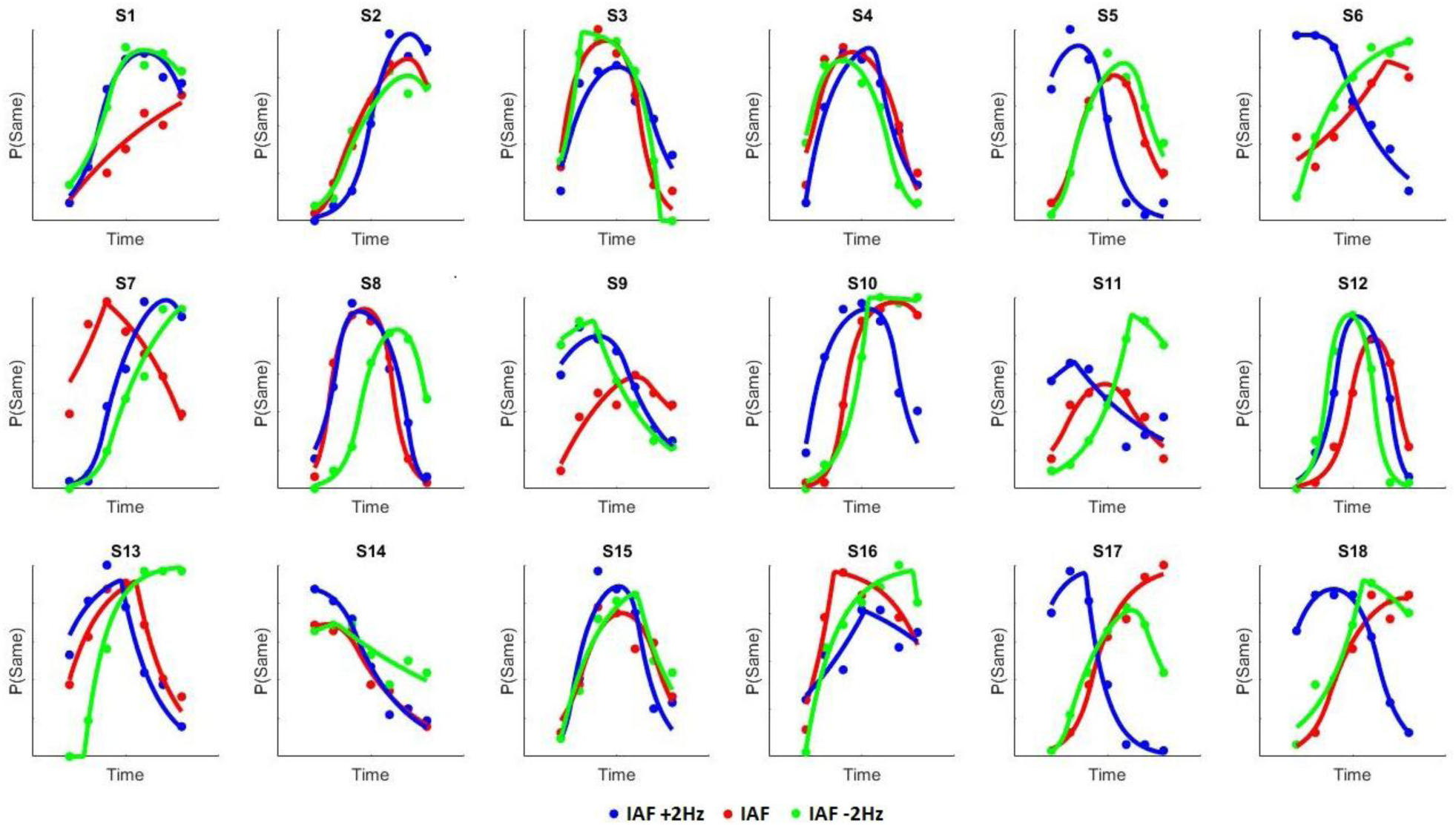
Individual psychometric functions for each of the stimulation conditions. Each point represents proportion of responding “same” for that tested interval.

**Figure 5.**
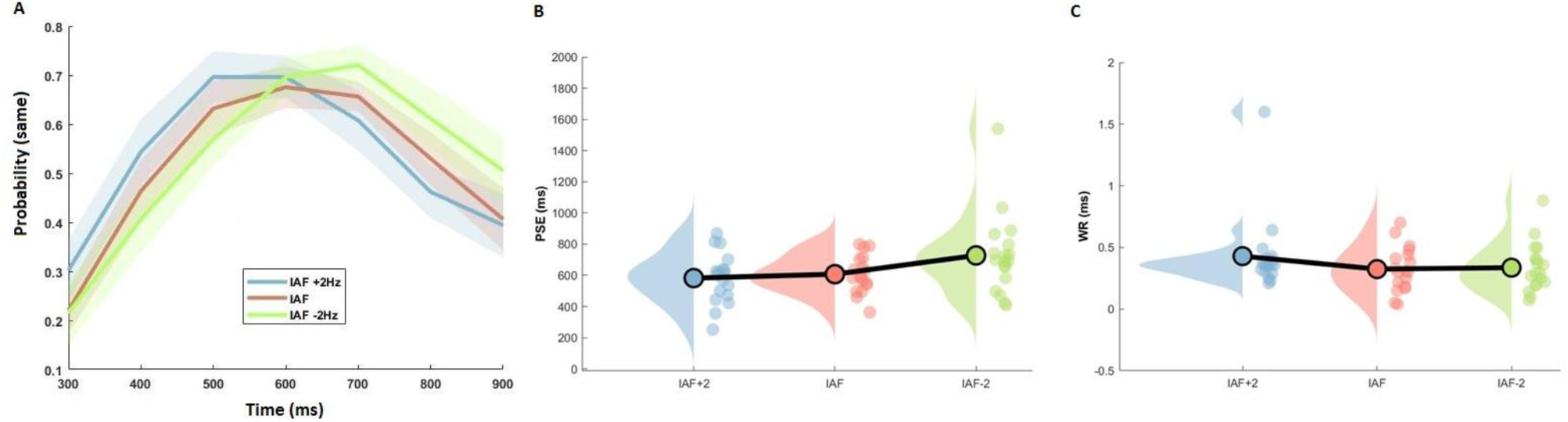
Behavioral results of tACS. **A)** Mean temporal generalization gradients for each of the stimulation conditions. Each point represents the average proportion of responding “same” for that tested interval. Subjects exhibited a leftward shift for IAF+2Hz and a rightward shift for IAF-2Hz, relative to IAF tACS. Shaded regions represent standard error. **B)** Raincloud plots of point of subjective equality (PSE) values across all three conditions exhibited the same pattern as in the average proportion data, with a linear shift in PSE for slower-to-faster tACS. **C)** Same plot at in (B), but for Weber ratio (WR) values, exhibiting no significant shifts across conditions.

Given the above findings, we further investigated if IAF interacted with the observed effect. To quantify this, we calculated the slope value of a linear regression for each subject between the PSE values and the individually-determined levels in stimulation frequency. Accordingly, a negative slope value indicates a shift towards longer/shorter perceived intervals with faster/slower frequencies of stimulation. When correlated with the baseline IAF values, we observed a positive correlation; this effect was quantified by both a Pearson [*r*(17)=0.523, *p* = 0.026] and Spearman [rho(17) = 0.481, *p* = 0.043] correlations and confirmed using 10,000 bootstrap samples for each [Pearson 95% confidence interval: 0.167 – 0.75; Spearman: 0.019 – 0.785]. As a further measure, we removed two subjects that could have overly driven the effect (Figure 6). The Pearson correlation remained significant (0.54, *p* = 0.025), whereas the Spearman became only marginally significant (0.47, *p* = 0.057). Thus, while we suggest the effect may have been influenced by a strong effect in two subjects, we note that even without these two the same relationship is observed.

**Figure 6.**
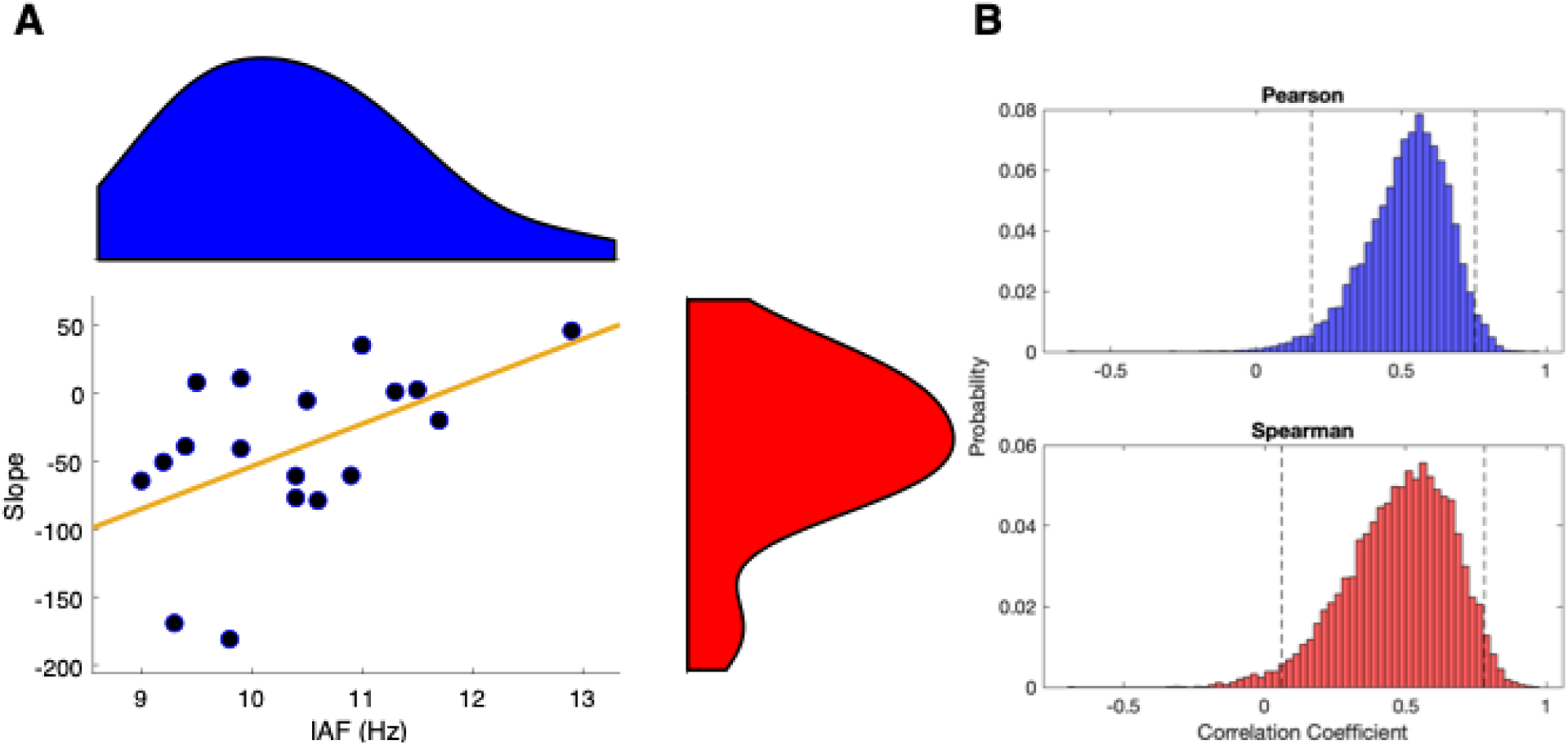
Correlation between baseline IAF and impact of tACS. **A)** Individual IAF values are plotted against the slope of a linear regression of PSE values across stimulation levels; a negative slope indicates that faster/slower tACS increased/decreased estimates of duration. Shaded regions indicate density estimates. A positive correlation was observed, indicating that subjects with a slower baseline IAF exhibited a larger impact of tACS. The effects remained largely significant with the removal of two outliers (see text). **B)** Bootstrapped distributions of Pearson and Spearman correlation coefficients; dashed lines indicate 95% confidence intervals for each distribution.

## Discussion

The present study aimed to investigate the link between modification of IAF and variation in perceived duration. In particular we hypothesized that applying tACS at IAF or driving IAF toward slower or faster oscillation rates would result in either under- or over-temporal estimation if IAF are involved in perceived duration, or in less- or more-precise estimation if involved in temporal resolution. We note that both possibilities could also have occurred, in which case under- or over-estimations would have become less- or more-precise. Our results confirm the first hypothesis by indicating that tACS applied at IAF ±2Hz produced a shift in the psychometric function consistent with a variation in perceived duration, with no change in the precision of temporal estimates.

The link between neural oscillations in the alpha range (8–12Hz) and temporal processing has a long history (Anliker, 1963; Wiener & Kanai, 2016). Several researchers suggested that α may contribute to defining psychological moments with the hypothesis that one α cycle (∼100 ms) would form the unit of subjective time, i.e., the psychological moment. Here, we extend previous findings by showing the intraindividual modification of IAF results in modified perceived duration. Importantly, we confirmed that the observed modification induced by tACS was specific for perceived duration and not on temporal variability, confirming a causal link between IAF and perceived duration. It is assumed that the speed of internal clock can be modulated by the speed of cortical oscillators, and in particular the clock speed could be represented by the alpha band regime (Treisman et al., 1990, 1994). Here we confirm the relationship between IAF and fluctuations in subjective timing. Further, our results demonstrate that baseline IAF rate contributes to the strength of the effect of tACS; subjects with a slower baseline IAF showed a greater influence of tACS than those with a faster IAF. This finding suggests an upper limit for speeding up alpha oscillations, such that, beyond a particular frequency, the visual system is unable to accommodate a faster clock signal.

Our results are consistent with previous studies that observed variation in perceived duration with transcranial Random Noise Stimulation (tRNS; Mioni, Grondin, Mapelli, & Stablum, 2018) compared to variation in temporal variability with transcranial Direct Currect Stimulation (tDCS; Mioni, Grondin, Forgione, Fracasso, Mapelli, & Stablum, 2016, Vicario, Martino, & Koch, 2013). tACS and tRNS are not intended to excite or inhibit cortical activity monotonously as it is for tDCS. Instead, the main goal of tACS is to influence brain oscillations (Woods et al., 2016). tRNS is similar to tACS in that it uses an alternating current; however, instead of stimulating at a fixed frequency throughout the stimulation period, tRNS alternates at a random frequency and amplitude within a specific range. As such, tACS provides the opportunity to test targeted hypotheses regarding specific frequency bands. We note that our results differ from another recent tACS study of time perception (Wiener et al., 2018) in which beta (20Hz), but not alpha stimulation induced a change in temporal perception. However, we further note that 1) stimulation in that study was localized to the supplementary motor area (SMA), 2) the study employed a temporal bisection task, with very different task demands and response mapping, and 3) the effect of beta oscillations was hypothesized to be a result of altering the memory, rather than perception, of perceived intervals, a finding consistent with beta’s potential role in timing (Mendoza et al., 2018). In particular, we also note that this previous study did not titrate stimulation frequency to individual values. This last point bears emphasizing: if we had stimulated all subjects at the same frequency within the alpha band (i.e. 10Hz) in the present study, no effect would have been observed, with some subjects speeding up and others slowing down.

Broadly, alpha oscillations have been associated with a variety of perceptual and cognitive processes, including attentional state and focus (Klimsch, 2012), perceptual awareness (Matthewson et al., 2009), problem solving (Fink & Neubauer, 2006), and working memory (Bonnefond & Jensen, 2012). Occipital alpha, which our experiment focuses on, has been hypothesized to underlie phasic transmission from the lateral geniculate nucleus to striate and extrastriate cortex (Bollimunta et al 2011; Vijayan & Kopell, 2012). Beyond the visual cortex, alpha oscillations have been linked with theta (∼5-7Hz) rhythms in coordinated patterns to prefrontal regions (Zhang et al., 2018). The function of this coordination has been associated with perceptual “sampling” of visual information, which may be probed via time-based illusions or behavioral cycles (van Rullen, 2016; Brüers & van Rullen, 2017; Tommassini et al., 2017). We here suggest, additionally, that this sample rate mediates the individual perception of duration. Further work will be necessary to determine how tACS specifically alters IAF and other EEG indices. Notably, tRNS, which also can shift perceived duration, has been associated with a frequency-specific increase in resting theta oscillations (van Doren et al., 2014) despite the broad, high range of frequencies employed. As theta oscillations are also involved in time perception (Kononowicz & van Rijn, 2015; Zold & Shuler, 2015; Wiener & Kanai, 2016), and are functionally coupled with alpha oscillations, it is possible that stimulation-specific effects on perceived, rather than remembered duration are the result of interactions between these two frequency bands, as tested by each stimulation technique. Consistent with this idea, recent evidence suggests that alpha-theta interactions underlie an intermixing of phasic sampling (alpha) and attentional focus (theta) such that each are necessary for gating of sensory information (Fiebelkorn et al., 2018; Helfrich et al., 2018).

Our results are also informative to neurobiological models of time perception. Recent debate exists in the field regarding the modal-specificity of duration processing, as to whether or not time depends on the modality in which it is perceived (Ivry & Schlerf, 2008; van Wassenhove, 2009; Merchant et al., 2013). Our results suggest that time is a modal feature, that may be altered by manipulating oscillatory cycling within the visual cortex, however, we note that our findings are limited in that the intervals tested were 1) sub-second, and 2) visually demarcated. Yet, to the former point, recent research suggests that visually-induced time dilation effects exist at both sub and supra-second durations (Wearden et al., 2014; Shima et al., 2016; Herbst et al., 2013), and both may be linked to occipital alpha (Hashimoto & Yotsumoto, 2018); to the latter, if timing is modal-specific, then an auditory version of our task would have resulted in a null effect. Further work will need to dissociate between the role of modality and interval length on the impact of tACS stimulation.

Overall, our results provide the first evidence that we are aware of for a causal link between individual alpha levels and the rate of the internal clock in time estimation. We further suggest that previous studies which could not account for a strong link between alpha and time estimation did not examine individual differences in alpha that may have accounted for heterogeneous results between subjects. Our results suggest a way forward for further studies to dissect an alpha-driven clock signal for the perception of duration.

## Acknowledgements

The information in this manuscript and the manuscript itself has never been published either electronically or in print. GM was suppsorted by “Iniziative di Cooperazione Universitaria 2018” sponsored by University of Padova. This work was carried out within the scope of the project “use-inspired basic research”, for which the Department of General Psychology of the University of Padova has been recognized as “Dipartimento di Eccellenza” by the Ministry of University and Research.

**The authors declare no competing financial interests**

